# Accuracy of a battery-powered portable capnometer in neonates

**DOI:** 10.1101/2020.06.05.136143

**Authors:** Eiji Hirakawa, Satoshi Ibara

## Abstract

End-tidal CO_2_ measurement (EtCO_2_) is useful for confirmation of successful tracheal intubation and ensuring adequate ventilation. There are two types of EtCO_2_ detector, i.e., single-use-only devices and capnometers. Although portable capnometers are widely used for resuscitation, there have been no reports regarding their clinical utility in neonates. Here, the correspondence between EtCO_2_ level determined using a battery-powered portable capnometer and arterial CO_2_ (PaCO_2_) was investigated using paired data obtained simultaneously from 26 neonates weighing 1262 ± 589 g at examination on mechanical ventilation. EtCO_2_ level and PaCO_2_ showed a strong correlation (r = 0.839, *P* < 0.001), and the correlation equation was: EtCO_2_ = 0.8 × PaCO_2_ + 1.1. Therefore, EtCO_2_ readings obtained with a battery-powered portable capnometer were likely to be underestimated. This became more pronounced with decreasing infant body weight at examination as the net difference in measurements of PaCO_2_ and EtCO_2_ was significantly positively correlated with infant body weight at examination (r = 0.451, *P* < 0.001). The observations presented here may be helpful in the use of battery-powered portable capnometers in neonates requiring controlled ventilation with tracheal intubation.

## Introduction

Measurement of carbon dioxide (CO_2_) using a CO_2_ detector is a useful method for confirming airway clearance [1]. However, the results obtained using a nasal CO_2_ detector are qualitative rather than quantitative. It is important to control CO_2_ in premature infants, especially during the acute phase, such as resuscitation, during transportation from the delivery room to the neonatal intensive care unit (NICU). As patients are on a ventilator in the NICU, the initial inspiratory pressure and respiratory rate are set automatically. However, during transportation after birth or resuscitation, ventilation is manually supported. The neonatal resuscitation program (NRP) currently doesn’t mention about monitoring CO2 due to lack of data at resuscitation, and continuous monitoring of EtCO2 during resuscitation of asphyxiated term lamb have been reported [2]. It is important to visually continuously confirm the EtCO_2_ using a capnometer, which helps the respiratory therapist to determine the appropriate initial settings when switching to a ventilator. The capnometer, which measures CO_2_ expired from the lung alveoli as end-tidal CO_2_ (EtCO_2_), is generally installed in anesthesia machines and is used to identify early airway incidents and to ensure adequate ventilation in the operating room, out of hospital, and during transport of critically ill patients [3, 4, 5].

However, as conventional capnometer devices are heavy, expensive, and of the built-in type, it is challenging to monitor EtCO_2_ in patients at the bedside or outside the hospital [4, 6]. Neonatal resuscitation is usually conducted in the delivery room or operation room, after which the infant is transported to the NICU. Therefore, there is a need for a portable capnometer that can be used easily at resuscitation or during transportation in emergency, delivery, and operation rooms. The EMMA™ capnograph (EMMA, Masimo, Danderyd, Sweden) is small, light (5.2 × 3.9 × 3.9 cm and 59.5 g with batteries), is readily portable, does not require routine calibration before use, can be warmed up in a short time, and can be used for continuous monitoring of EtCO_2_ [7]. In addition, this is the only portable capnometer currently available with 1-mL dead space for neonates.

Respiratory function includes oxygenation of blood and expiration of CO_2_. Premature neonates often have difficulties in respiration and require tracheal intubation immediately after birth for controlled ventilation. SpO_2_ monitoring is recommended during resuscitation and transportation of neonates with respiratory failure to the NICU [8]. However, SpO_2_ monitoring alone without EtCO_2_ monitoring is insufficient as SpO_2_ is only an indicator of the ratio of combined O_2_ to hemoglobin. The SpO2 monitor provides no information about CO_2_ levels and is not useful to estimate respiratory acidosis caused by hypercapnia due to inadequate respiration. In neonates, although hypercapnia has been recognized as permissive hypercapnia to protect the lungs, hypercapnia and subsequent respiratory acidosis, which occur after birth, may exacerbate pulmonary hypertension and make it persistent [9]. On the other hand, hypocapnia is related to brain injury [10]. Hyperventilation with subsequent hypocapnia was a widely practiced strategy in the past [11]. In the 1980s, hyperventilation was the only method available for the treatment of persistent pulmonary hypertension (PPHN), as inhalation of nitric oxide (NO) and/or extracorporeal membrane oxygenation (ECMO) were not commonly accepted treatment options. However, artificially initiated hypocapnia is now recognized as a major risk factor for intracranial intraventricular hemorrhage (IVH) or periventricular leukomalacia (PVL) [12]. Hypocapnia causes brain vasoconstriction, and thus reduces cerebral blood flow (CBF) [11].

The brain vasculature at watershed areas is particularly underdeveloped and vulnerable to hypoperfusion in premature infants at around 32 weeks of gestation, and PVL can occur in this area. IVH is caused by multiple reasons, one of which is hypocapnia [11, 12, 13]. Severe IVH is related to neurodevelopmental delay, and 48% of cases of IVH are observed within the first 6 hours of life. Therefore, strict control of blood CO_2_ levels in the acute phase is required [13, 14].

A portable device to monitor EtCO_2_ is expected to facilitate continuous monitoring of blood CO_2_ levels during neonatal resuscitation and/or transportation to the NICU. To our knowledge, however, there have been no studies focusing on the accuracy of the EMMA capnograph in neonates. This study was performed to assess the accuracy of the EMMA capnograph in neonates on mechanical ventilation by comparing ETCO_2_ measured using this device with arterial carbon dioxide (PaCO_2_) level.

## Materials and Methods

This study was approval by the institutional review board (IRB) of Kagoshima City Hospital (KCH) (20150903). Neonates admitted to the NICU of KCH between July 2014 and February 2015 and requiring controlled ventilation were eligible for inclusion in the study. The parents of all patients provided written informed consent before participation in this study. Two mainstream capnographs are used for patients on mechanical ventilation in KCH, EMMA and CapONE™ (Nihon Kohden, Tokyo). Sidestream capnography is not used. The EMMA was placed proximal to the endotracheal tube using a neonatal adaptor with 1.0-mL dead space. PaCO_2_ levels were measured using 100-μL arterial blood sample, and EtCO_2_ levels were recorded at the same time as sampling as a number without a decimal point. Each participant underwent EtCO_2_ measurement at least once a day.

Participants on mechanical ventilation required repeated EtCO_2_ measurement during their hospital stay when arterial blood samples were collected. PaCO_2_ was measured using an ABL 800™ (Radiometer, Copenhagen, Denmark). All measurements were recorded with the patient in the supine position. Blood gas analysis was performed when ordered by the attending neonatologist depending on the condition of the patient.

Statistical analyses were performed using the JMP13© statistical software package (SAS, Cary, NC). The Wilcoxon/Kruskal–Wallis method was used for analysis of continuous variables. Fisher’s exact probability test was used for analysis of categorical variables. Regression analysis was performed. In all analyses, *P* < 0.05 was considered statistically significant.

## Results

The study population consisted of 26 newborn infants (Table 1). The median (minimum − maximum) gestational age at birth was 27 (22 – 38) weeks and median birth weight was 881 (420 – 2730) g. Tracheal tubes were selected according to birth weight as follows: < 1000 g, Fr. 2.5; 1000 – 1800 g, Fr. 3.0; and > 1800 g, Fr. 3.5. At the point of measurement, the ventilation mode was synchronized intermittent mechanical ventilation (SIMV) without volume guarantee and the leakage displayed on the ventilator was 0% in all patients. Simultaneous EMMA EtCO_2_ and PaCO_2_ measurements were performed 63 times in the 26 infants (median, three times per infant). Median values of corrected age, body weight, and tidal volume of those measured with EMMA were 30 (22 – 38) weeks of gestation, 1151 (420 – 2730) g, and 5.6 (3.3 – 7.4) mL/kg, respectively (Table 1). No adverse events associated with use of the EMMA capnograph occurred in the 26 infants.

**Table 1.**
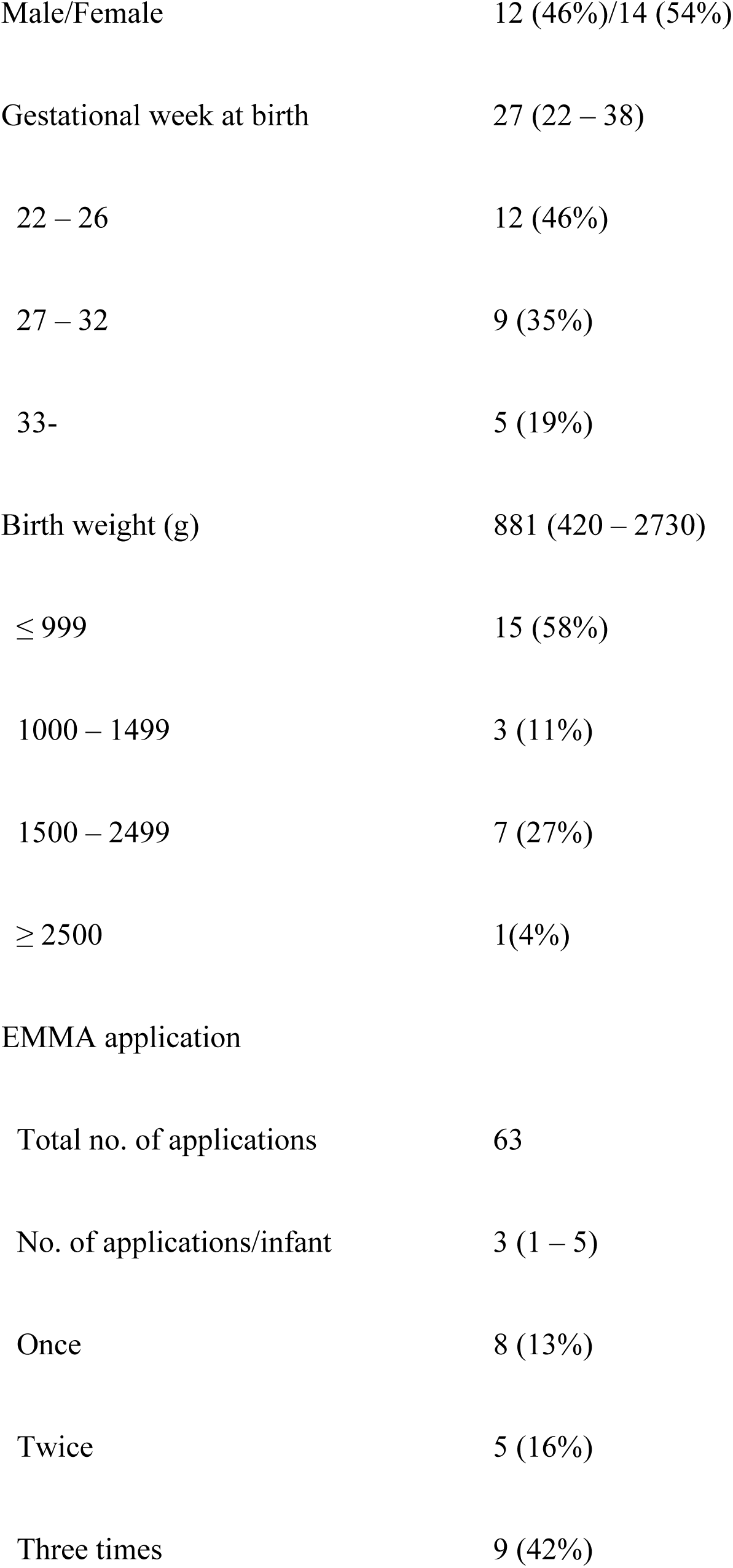
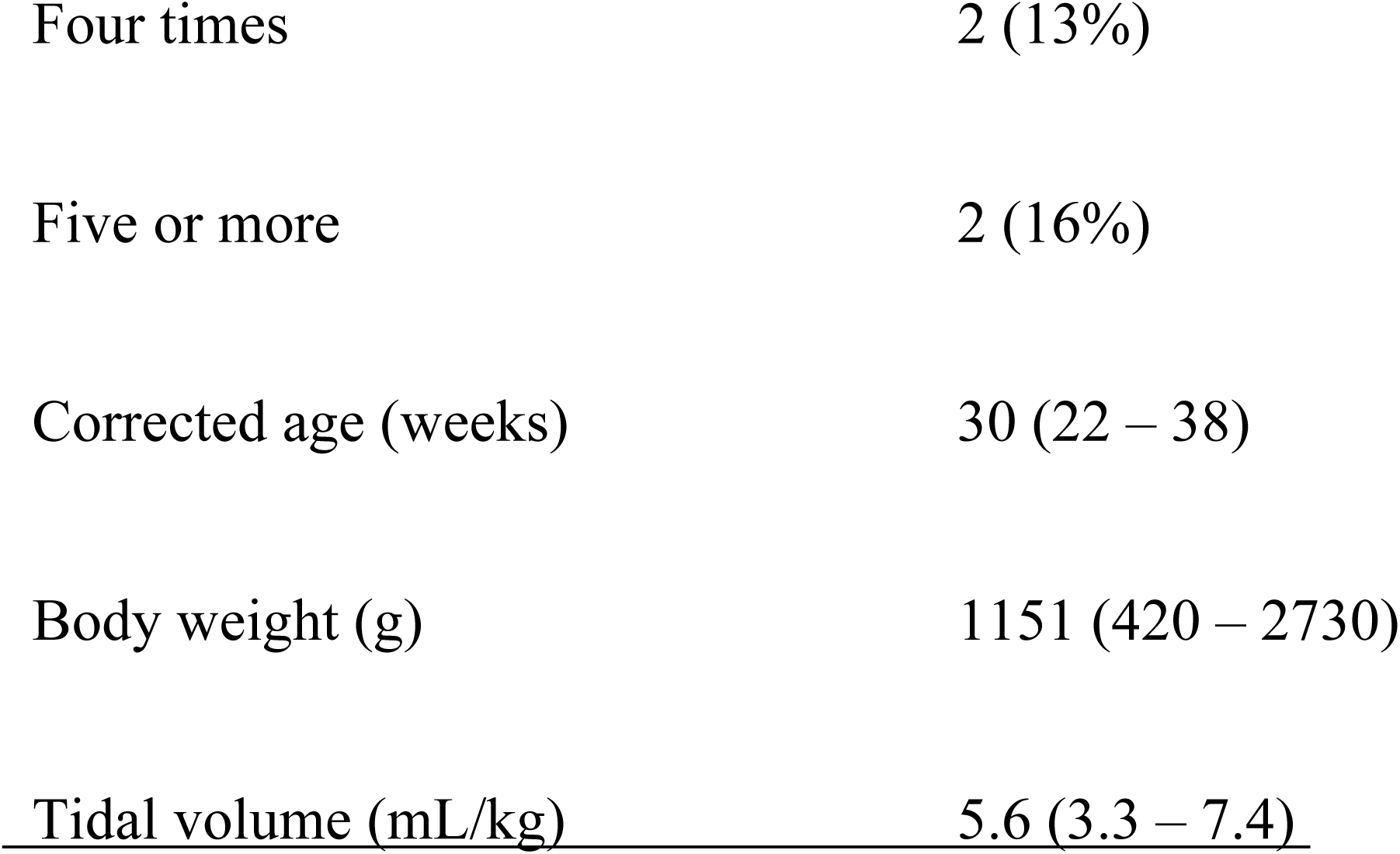
Demographic characteristics of the 26 patients

### Correlation between EtCO_2_ and PaCO_2_

As shown in Fig. 1, the majority of EtCO_2_ measurements were lower than those of PaCO_2_. However, the correlation coefficient (r) was 0.839 and the correlation equation was as follows for 63 paired data: EtCO_2_ = 0.80 × PaCO_2_ + 1.17. Thus, the overall EtCO_2_ measurements were approximately 20% lower than PaCO_2_ measurements.

**Figure 1.**
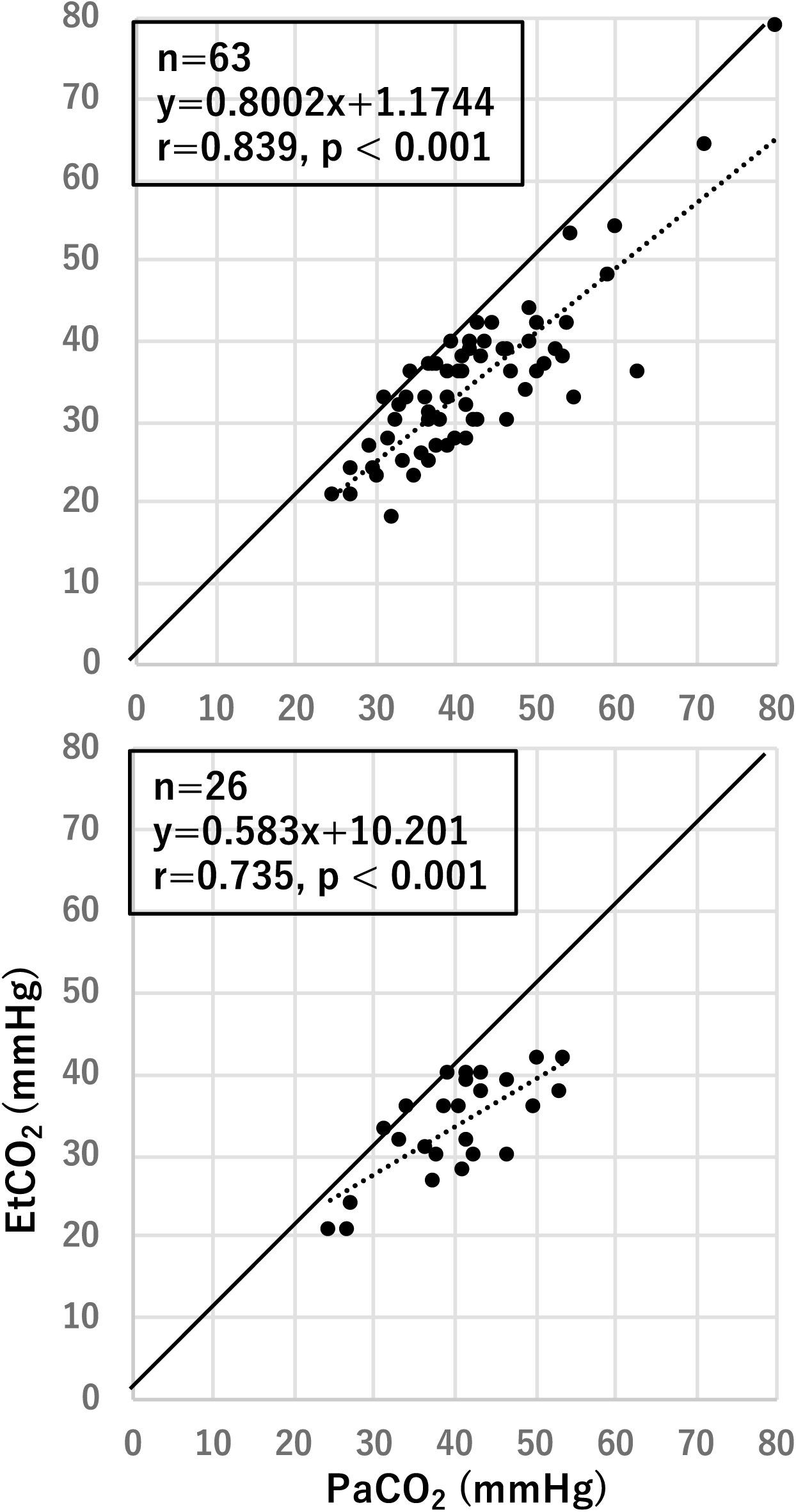
Correlation between PaCO_2_ and EtCO_2_ PaCO_2_ and EtCO_2_ were determined simultaneously 63 times in the 26 infants.

### Effects of infant body weight on EtCO_2_ measurements

The body weights of the infants showed no significant correlation with PaCO_2_ measurements, but were significantly positively correlated with EtCO_2_ measurements (Fig. 2, upper and middle panels). As most EtCO_2_ measurements were lower than those of PaCO_2_ (Fig. 1), these results suggested that EtCO_2_ measurements were more likely to be underestimated in infants with lower body weight. This was the case, as shown in the lower panel of Fig. 2, where net differences between measurements of PaCO_2_ and EtCO_2_ increased as infant body weight decreased. However, the net differences were not significantly correlated with tidal volume (mL/kg) at the time of examination.

**Figure 2.**
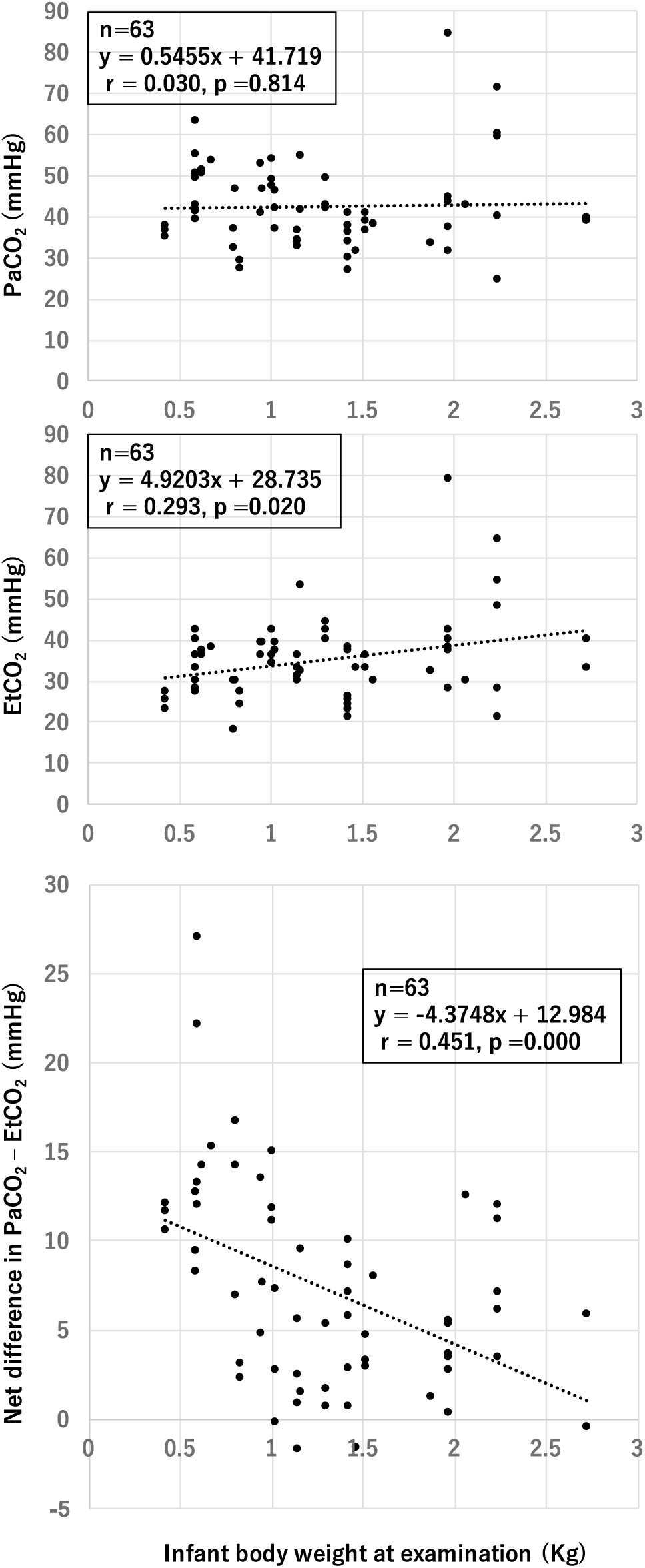
Effects of infant body weight at examination on PaCO_2_, EtCO_2_, and net difference in measurements of PaCO_2_ and EtCO_2_

## Discussion

In this study, EtCO_2_ measured using an EMMA capnograph in neonates was strongly correlated with PaCO_2_. There have been no previous reports regarding the correlation between EtCO_2_ measured using an EMMA capnograph and PaCO_2_ in neonates. This is the first report of the application of this device to neonates, including extremely infants with low birth weight. As shown in Fig.1, correlations between EtCO_2_ and PaCO_2_ were observed over wide ranges; PaCO_2_ [24.5 – 84.5] and EtCO_2_ [18 – 79]. These observations also showed that the EMMA capnograph correctly followed PaCO_2_ over a wide range. However, EtCO_2_ was approximately 20% lower than PaCO_2_. The EMMA capnograph underestimated blood levels of CO_2_ by 20% compared with PaCO_2_. Singh et al. reported a correlation between PaCO_2_ and EtCO_2_ following surfactant therapy [15]. Hyaline membrane disease (HMD) is also one possible explanation for the difference [16]. However, no infants with HMD were included in the present study. Our data showed that the net difference was larger in infants with lower birth weight, meaning that a greater tidal volume was required for accuracy of EtCO_2_ measurement. EtCO_2_ was measured with a sensor set at the proximal end of the tracheal tube, which was also the end of the entire ventilation circuit. Therefore, tidal volume (mL) may affect EtCO_2_ depending on the level of flow at the sensor. In management of mechanical ventilation, tidal volume per weight (TV mL/kg) is a score that ensures that he ventilator settings are adequate for the patient. The normal range of TV mL/kg in neonates is 4 – 6 mL/kg. Obviously, the net tidal volume (mL) is smaller in infants with lower birth weight. In this study, the median body weight and median TV mL/kg at the time of examination were 1151 (420 – 2730) g and 5.6 (3.3 – 7.4) mL/kg, respectively. TV mL/kg was adequate for each body weight. Low tidal volume may have been responsible for the observed net differences between PaCO_2_ and EtCO_2_. The differences may also have been due to differences in the sizes of tracheal tubes. Use of a tracheal tube of inadequate size may affect tidal volume, because the tracheal tubes used in neonates do not have a cuff, and therefore tidal volume may be affected by air leakage. However, air leakage displayed on the ventilator monitors was 0% at each examination in this study.

## Conclusion

The results of this study indicated that EtCO_2_ measured using an EMMA capnograph was positively correlated to PaCO_2_ even in preterm neonates. Thus, this device may be applicable even in preterm neonates. As shown in the correlation equation, EtCO_2_ determined with the EMMA capnograph was approximately 20% lower than PaCO_2_. Therefore, this underestimation must be taken into account when assessing blood CO_2_ level using this device.

## Author contributions

E.H., and S.I. contributed to the study design and writing the manuscript.

All authors have seen and approved the final version of this manuscript.

The authors declare no conflict of interest.

